# A genome-wide atlas of antibiotic susceptibility targets and pathways to tolerance

**DOI:** 10.1101/2022.01.26.477867

**Authors:** Dmitry Leshchiner, Federico Rosconi, Bharathi Sundaresh, Emily Rudmann, Luisa Maria Nieto Ramirez, Andrew T. Nishimoto, Stephen J. Wood, Bimal Jana, Noemí Buján, Kaiching Li, Jianmin Gao, Matthew Frank, Stephanie M. Reeve, Richard E. Lee, Charles O. Rock, Jason W. Rosch, Tim van Opijnen

**Affiliations:** Boston College, Biology Department, Chestnut Hill, MA, 02467, USA; Department of Infectious Diseases, St. Jude Children’s Research Hospital, Memphis, TN 38105, USA; Boston College, Chemistry Department, Chestnut Hill, MA, 02467, USA; Department of Chemical Biology and Therapeutics, St. Jude Children’s Research Hospital, Memphis, TN 38105, USA

## Abstract

Detailed knowledge on how bacteria evade antibiotics and eventually develop resistance could open avenues for novel therapeutics and diagnostics. It is thereby key to develop a comprehensive genome-wide understanding of how bacteria process antibiotic stress, and how modulation of the involved processes affects their ability to overcome said stress. Here we undertake a comprehensive genetic analysis of how the major human pathogen *Streptococcus pneumoniae* responds to 20 antibiotics. We built a genome-wide atlas of drug susceptibility determinants and generate a genetic interaction network that connects cellular processes and genes of unknown function, which we show can be used as therapeutic targets. Pathway analysis reveals a genome-wide “tolerome”, defined by cellular processes that can make a bacterium less susceptible, and often tolerant, in an antibiotic specific manner. Importantly, modulation of these processes confers fitness benefits during active infections under antibiotic selection. Moreover, screening of sequenced clinical isolates demonstrates that mutations in tolerome genes readily evolve and are frequently associated with resistant strains, indicating such mutations may be an important harbinger for the emergence of antibiotic resistance.

## INTRODUCTION

The emergence of antibiotic resistance in bacterial pathogens is a continuously developing complex problem that is only solvable if besides new drugs we also learn to understand the exact (genetic) processes that enable resistance. For instance, new antibiotics and treatment strategies are key to retain the ability to treat resistant infections. However, a comprehensive understanding of how and under which conditions resistance emerges, which genes and pathways contribute to drug sensitivity, and how resistance may be prevented or even taken advantage of, are equally important, as it could make treatments more focused and possibly less dependent on new drugs. For many antibiotics we know which genomic changes can cause resistance. However, it is often not clear how we get there with respect to which evolutionary paths are taken and whether for instance tolerance or lowered drug sensitivity precedes resistance. Interestingly, clinical strains isolated during antibiotic treatment failure may lack known resistance markers and instead contain multiple changes that may have no clear or known role in resistance^1-5^. However, whether these changes play a role or not is often unclear because the distribution of changes that can affect a bacterium’s drug sensitivity are largely unknown^1-7^. Therefore, understanding which genes, pathways and processes can contribute to altered drug susceptibility, could help identify genomic changes that not only sensitize bacteria to certain drugs, but desensitize them and may thereby act as precursors for antibiotic escape and/or resistance development.

Resistance emerges primarily through drug target mutations blocking antibiotic lethal action, upregulation of efflux pumps, and the acquisition of drug inactivating enzymes ^7-13^. Importantly, an antibiotic’s effects go far beyond the interaction with its direct target. We, and others, have shown that when a bacterium is challenged by an antibiotic, the imposing stress can expand throughout the bacterium and affect and demand the involvement of many different processes ^6,14-17^. For instance, while fluoroquinolones like ciprofloxacin inhibit DNA replication by targeting gyrase and/or topoisomerase, this also triggers double stranded breaks requiring the involvement of DNA repair mechanisms, which in turn requires nucleotide and energy metabolism. Antibiotics can thereby trigger a stress cascade, that with mounting stress increasingly reverberates through the organismal network, until the accumulating stress passes a threshold at which point the organism succumbs to the pressure ^15,17^. This explains why mutations in genes or pathways involved in dealing with the downstream (indirect) effects of antibiotic exposure can often make a bacterium more sensitive to a specific antibiotic. Indeed, we have shown for *Streptococcus pneumoniae* and *Acinetobacter baumannii* that, for instance, targeting DNA repair makes bacteria more susceptible to fluoroquinolones^6,16,18^, or targeting the Rod-system and/or Divisome makes *A. baumannii* more sensitive to cell wall synthesis inhibitors (CWSIs)^6^. This means that downstream genes, pathways and processes can be used as new targets or drug potentiators, either by themselves or in combination with others^6,14^. Moreover, in most bacteria, as in any other organism, the majority of genes are of unknown function, it is unclear what role they play in a specific process and/or pathway, or how they are connected within the organismal genomic network. Thus, besides solving gene-function, mapping-out which genes, pathways and processes are involved in dealing with and overcoming antibiotic-stress, and how they interact with each other, can provide key insights into uncovering new drug targets, or for instance rational combination strategies^6^.

While identifying off-target genes and pathways that increase drug sensitivity may thus be useful, it is possible that changes in associated processes could, in contrast, just as well reduce the experienced antibiotic stress. Such changes would thereby decrease antibiotic sensitivity and could possibly function as precursors to the emergence of resistance. A possible example of this is the induction of tolerance and/or persistence, where a small proportion of bacterial cells in the population upon exposure to high (transient) concentrations of antibiotics, are induced into a cell state that enables them to tolerate this treatment. Cell states associated with tolerance include cell dormancy, slow growth, transient expression of efflux pumps, and induction of stress response pathways ^19,20,21,22^. However, the mechanistic underpinnings of tolerance and decreased antibiotic sensitivity remain largely undefined and possibly differ between bacterial species and vary among antibiotics^23^. Moreover, specific mutations can (dramatically) increase the fraction of the surviving population ^24-26^, indicating these tolerant phenotypes have a genetic basis. Lastly, since clinical isolates often carry mutations located outside well-characterized drug targets ^1-5,27,28^, they could thus be composed of variants with different antibiotic sensitivities. Consequently, such variants with decreased antibiotic sensitivity could enable antibiotic escape, and/or enable multi-step high-level resistance mutations to evolve as they are given an extended opportunity to emerge^21,29-32^. Variants with decreased antibiotic sensitivity may thereby play an important role in antibiotic treatment failure ^5,33,34^. However, the breadth of possible genetic alterations that can trigger tolerance and/or decrease antibiotic sensitivity are largely unknown, making it unclear how often and probable it is that such variants arise.

In this study we use Tn-Seq in *S. pneumoniae* exposed to 20 antibiotics, 17 additional environments, and two *in vivo* infection conditions, to generate a genome-wide atlas of drug susceptibility determinants and build a genome-wide interaction network that connects cellular processes and genes of unknown function. We explore several interactions as new leads for gene function, while we show that specific interactions can be used to guide the identification of targets for new antimicrobial strategies. We highlight one such novel target in the membrane, by successfully developing a combinatorial antibiotic-antibody strategy that significantly reduces the bacterial load during an acute mouse lung infection. Furthermore, detailed mapping of antibiotic sensitivity data to pathways and genes with known function suggests a genome-wide “tolerome” exists defined by a multitude of genomic changes to a wide variety of pathways and processes that can make the bacterium less susceptible, and often tolerant to specific antibiotics. We untangle some of the underlying genetic mechanisms and show that decreased susceptibility and/or tolerance can come from a variety of changes including those in (nucleotide) metabolism, (p)ppGpp and ATP synthesis, transcription and translation, as well as different types of transport. By further combining *in vivo*-infection-with antibiotic-Tn-Seq we predict and experimentally validate that many disruptions may retain their decreased antibiotic sensitivity phenotype *in vivo*, and thereby outcompete the wildtype in the presence of antibiotics. Moreover, by screening hundreds of clinical isolates we show that changes in tolerome genes readily evolve in human patients and are often associated with antibiotic resistance. Consequently, these data highlight the wide array of possibilities that can lead to lowered antibiotic sensitivity and/or tolerance and underscore the importance of understanding the genetics of variants with altered drug susceptibility.

## RESULTS

### A genome-wide view of antibiotic sensitivity

To obtain a genome-wide view of the genetic determinants that can modulate antibiotic stress in *S. pneumoniae*, Tn-Seq was employed in the presence of 20 antibiotics (ABXs), representing four classes and 9 different ABX groups (Fig. 1a). Six independent transposon libraries were generated and grown for approximately 8 generations in the absence and presence of an antibiotic at a concentration that reduces growth by approximately 30-50% (Supplementary Table 1). Tn-mutant frequencies are determined through Illumina sequencing from the beginning and end of the experiment with high reproducibility between libraries (R^2^ = 0.70-0.90; Supplementary Fig. 1) which is consistent with previous Tn-Seq experiments^6,15,16,18,35-38^. Combined with the population expansion during the experiment each mutant’s fitness (*W*) is calculated to represent their environment-specific relative growth rate ^6,18,35,39,40^. Each gene’s antibiotic-specific fitness is statistically compared to baseline fitness without ABXs, and is represented as *ΔW* (*W*_*ABX*_ - *W*_*noABX*_) and categorized as: **1)** Neutral, *ΔW* = 0, a mutant’s relative growth is similar in the absence and presence of an ABX; 2) Negative, *ΔW* < 0, a mutant’s fitness is significantly lower and thus grows relatively slower in the presence of an ABX; 3) Positive, *ΔW* > 0, a mutant’s fitness is significantly higher and thus grows relatively faster in the presence of an ABX. All antibiotics trigger both positive and negative growth effects (Fig. 1b, Supplementary Table 2), which are distributed across 22 different gene categories (Fig. 1c). Importantly, enrichment analysis shows there are multiple expected patterns, for instance genes involved in DNA-repair are enriched in the presence of fluoroquinolones; cell-wall, peptidoglycan and cell division genes are enriched in ß-lactams and glycopeptides; membrane integrity genes in lipopeptides; and transcription and translation in PSIs (Fig. 1d). Additionally, throughout the manuscript we validate a total of 49 predicted genotype x phenotype interactions, which indicates the Tn-Seq data is in line with previously shown accuracy^6,15,16,18,35-38^, and of high quality (Fig. 1e, Supplementary Table 8).

**Figure 1.**
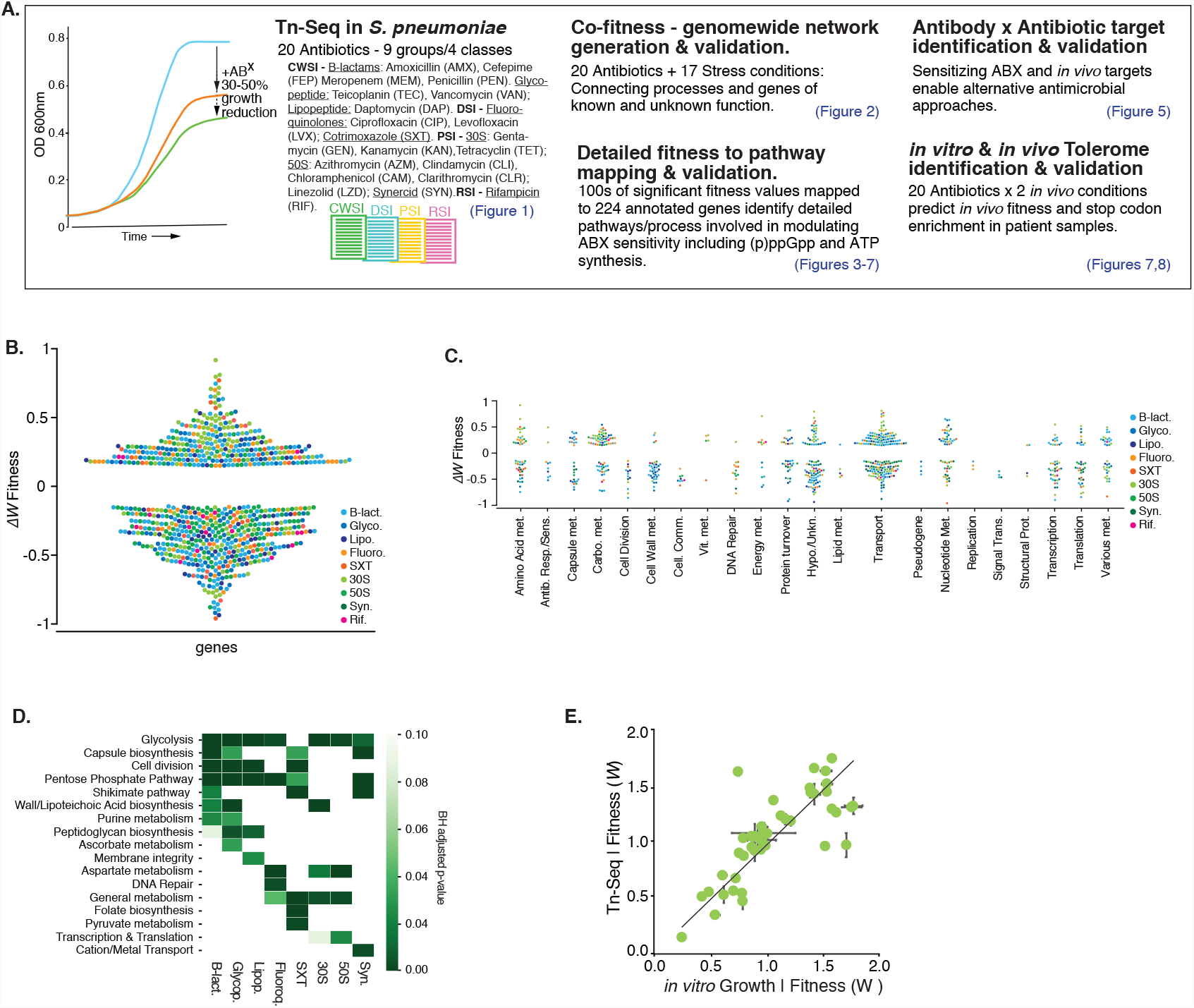
A genome-wide atlas of negative and positive fitness effects, highlights a multitude of processes that can modulate antibiotic susceptibility. **a**. Project setup and overview. Tn-Seq is performed with *S. pneumoniae* TIGR4, which is exposed to 20 antibiotics at a concentration that reduces growth by 30-50%. Genome-wide fitness is determined for each condition, suggesting a multitude of options exists to increase as well as decrease antibiotic sensitivity. A co-fitness network is constructed by adding Tn-Seq data from 17 additional conditions, which through a SAFE analysis highlights functional clusters, and connects known and unknown processes. The genome-wide atlas and network are used to develop an antibiotic-antibiotic combination strategy, and to map out the wide-ranging options that can lead to decreased antibiotic sensitivity *in vitro* and *in vivo* and that are associated with a higher rate of stop-codons in clinical samples. **b**. There are a large number of genetic options that can modulate antibiotic sensitivity; with significant increased (*Δ*W < −0.15) and decreased sensitivity (*Δ*W > 0.15) split over all antibiotics almost equally likely. **c**. Additionally, increased and decreased antibiotic sensitivity are distributed across a wide variety of functional categories. **d**. Enrichment analysis shows that some pathways/processes such as glycolysis are relatively often involved in modulating responses to antibiotics, while other processes are more specific. **e**. Validated growth experiments performed throughout the project highlight the Tn-Seq data is of high quality. S.E.M. bars are shown.

### Co-fitness interaction networks identify known and unknown genetic relationships

Screens such as Tn-Seq are geared towards highlighting the processes and genes that are important under a specific screening condition. With increasing conditions, genes acquire specific profiles, and those with similar fitness profiles can help reveal pathways and/or gene-clusters with similar and/or shared tasks. To extract such patterns, we build a correlation matrix based on each gene’s fitness-profile generated from 20 antibiotics and supplemented with previously collected Tn-Seq data from 17 additional non-antibiotic conditions^18^ (Supplementary Table 3). This results in a 1519×1519 gene matrix where positive correlations between genes come from shared phenotypes, while negative correlations come from opposing phenotypic responses under the same condition (Supplementary Table 4). By repeatedly hiding random parts of the data the stability and strength of each correlation is calculated and represented in a stability score (Supplementary Table 5). The correlation matrix and stability score are turned into a network, where each node is a gene, and each edge is a correlation coefficient above a threshold (>0.75), which combined with the stability score indicates the strength of the relationship between two genes. (Fig. 2a; Supplementary Table 6). Spatial Analysis of Functional Enrichment (SAFE)^41,42^ is used to define local neighborhoods within the network, i.e., areas enriched for a specific attribute (e.g., a pathway or functional category), which identifies multiple clusters that represent specific pathways and processes including purine metabolism, cell-wall metabolism, cell division and DNA repair (Fig. 2b; Supplementary Table 7). Moreover, the network contains gene-clusters of high connectivity identifying highly related genes including those within the same operon such as the *ami*-operon, an oligo-peptide transporter, the *dlt*-operon which decorates wall and lipoteichoic acids with d-alanine, and the *pst*-operon a phosphate transporter (Fig. 2c, I-III)). Besides identifying known relationships, the network also uncovers interaction clusters between genes with known and unknown interactions and function. Several such clusters are highlighted in Fig. 2c (IV-VIII), including genes involved in purine metabolism (further explored below), threonine metabolism, and in secretion of serine rich repeat proteins (SRRPs), which are important for biofilm formation and virulence^43^. Importantly, the identification of biologically relevant relationships among (clusters of) genes indicates the data is rich in known and new information.

**Figure 2.**
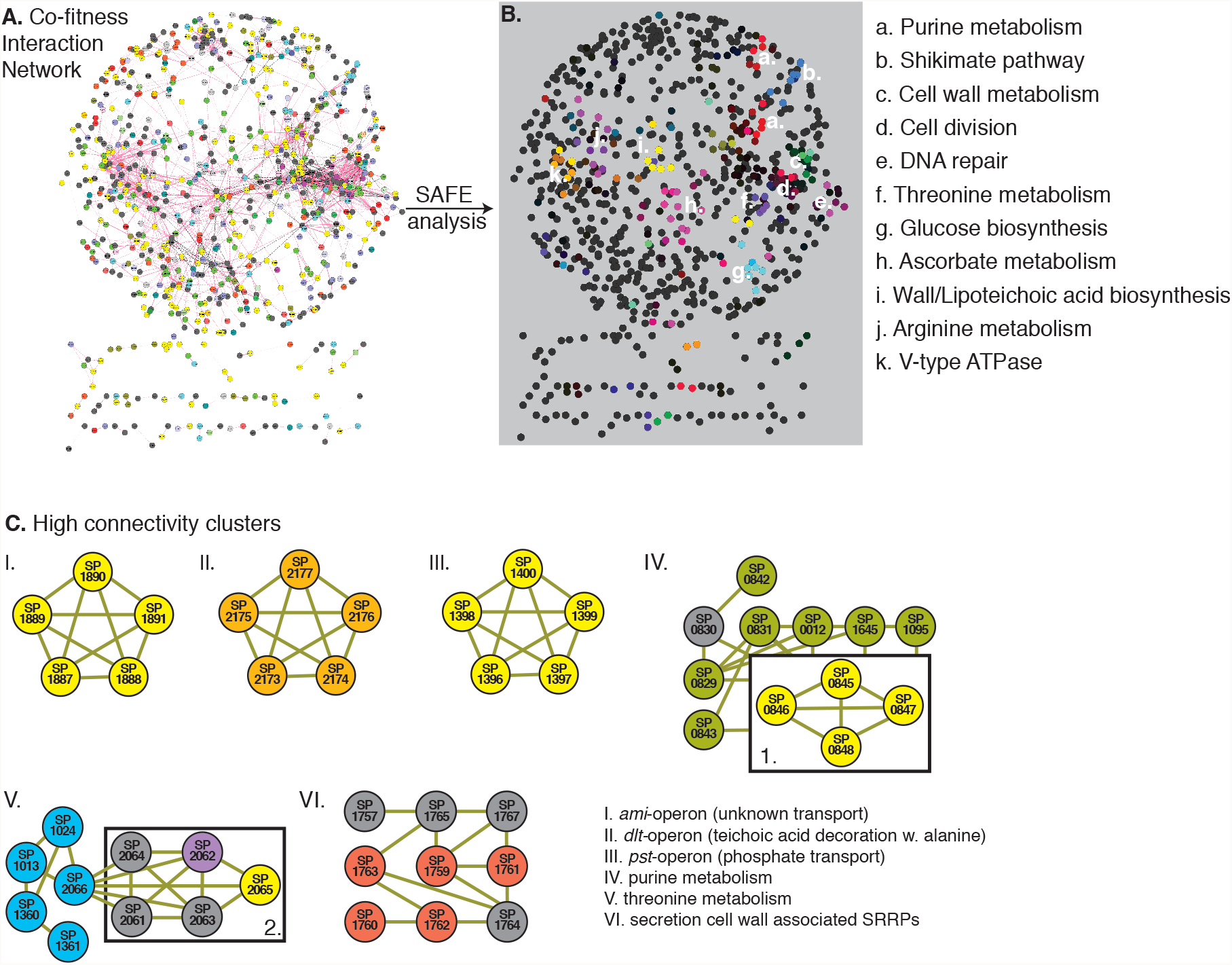
A co-fitness network identifies tight genetic clusters of known and unknown genes and processes. **a**. A 1519×1519 gene correlation matrix based on Tn-Seq data from 37 conditions generates a network with genes as nodes, and edges as interactions with a stability score and thresholded correlation >0.75. The network contains one large connected component and multiple smaller components placed underneath; **b**. A SAFE analysis identifies at least 11 clusters within the network that represent specific pathways and processes; **c**. The network contains highly connected clusters of smaller groups of genes for instance those within the same operon such as cluster I. the *ami*-operon, a putative oligo-peptide transporter; II. the *dlt*-operon which decorates wall and lipoteichoic acids with d-alanine; and III. the *pst*-operon a phosphate transporter. Several additional clusters are highlighted containing annotated and unannotated genes, connected through known and unknown interactions including cluster IV, which contains genes involved in purine metabolism (green nodes) and a putative deoxyribose transporter (yellow; boxed 1.); V. genes involved in threonine metabolism (blue) and several genes located as neighbors to SP_2066/*thrC* with unclear functions (boxed 2), including a regulator (SP_2062; purple) and a transporter (SP_2065; yellow); VI. genes involved in secretion of serine rich repeat proteins (SRRPs), which are important for biofilm formation and virulence, grey-nodes are unannotated genes.

### Detailed pathway mapping identifies a multitude of antibiotic susceptibility targets and pathways to tolerance

224 genes with a known annotation are present in the data that have at least one significant phenotype in response to an antibiotic, and which can be split over 21 functional groups according to a pathway or process they belong to (Fig. 3a). Each group is characterized by having multiple phenotypes that increase sensitivity in response to one or more antibiotics (negative phenotype), while each group, except for cell division, also has multiple phenotypes that decrease antibiotic sensitivity (Fig. 3a; positive phenotype). Moreover, each antibiotic group triggers both negative and positive effects (Fig. 3b). Where possible, the 21 functional groups are organized according to a pathway they belong to and/or relationships among genes and combined with their antibiotic susceptibility profile. This results in an antibiotic susceptibility atlas, which shows on a fine-grained scale, how inhibiting a pathway or process can affect sensitivity to an antibiotic (Fig. 3c and Supplementary Fig. 2 and 3). For instance, in the glycolysis-group, knocking out any of the three genes involved in forming the phosphotransferase (PTS)-system (SP_0282-SP_0284) that imports glucose to generate glucose-6-phosphate (G-6P), has a negative effect on fitness in the presence of 30S and 50S PSIs as well as Synercid (a synergistic combination of two PSIs), while it increases fitness in the presence of all CWSIs (ß-lactams, glycopeptides, and daptomycin) and fluoroquinolones. Also, the inhibition of/knocking out SP_0668 (*gki*, glucokinase), an enzyme that converts *α*-D-Glucose into G-6P, has a positive effect on fitness in all CWSIs and a negative effect in 30S PSIs. In contrast, inhibiting SP_1498 (*pgm*, phosphoglucomutase), the major interconversion enzyme of G-6P and G-1P, has a negative effect on fitness with all antibiotics (Fig. 3c). Within pyruvate metabolism, inhibiting lactate (SP_1220, *ldh*, L-lactate dehydrogenase), or acetaldehyde production (SP_2026, alcohol dehydrogenase) increases sensitivity to ß-lactams and glycopeptides and decreases sensitivity to 30S PSIs; inhibiting formate production (SP_0459 [*pfl*, formate acetyltransferase] and SP_1976 [*pflA*, pyruvate formate lyase activating enzyme]) decreases sensitivity to co-trimoxazole and 30S PSIs, while inhibiting acetyl-phosphate production (SP_0730, *spxB*, pyruvate oxidase) decreases sensitivity to ß-lactams, glycopeptides and co-trimoxazole. Within aspartate metabolism, interfering with SP_1068 (*ppc*, phosphoenolpyruvate carboxylase), which generates oxaloacetate from phosphoenolpyruvate (PEP), triggers a range of changes from increased sensitivity to ß-lactams, and glycopeptides, to decreased sensitivity to most other antibiotics, while the four genes involved in the production of threonine from L-aspartate (SP_0413 [aspartate kinase], SP_1013 [*asd*, aspartate semialdehyde dehydrogenase], SP_1360 [*thrB*, homoserine kinase], SP_1361 [*him*, homoserine dehydrogenase]) trigger decreased sensitivity to fluoroquinolones and 30S and 50S PSIs. In the shikimate pathway inhibiting the production of chorismate from PEP and erythrose-5-phosphate (through genes SP_1370 [*aroK*], SP_1371 [*aroA*], SP_1374 [*aroC*], SP_1375 [*aroB*], SP_1376 [*aroE*], SP_1377 [*aroD*]) leads to increased sensitivity to ß-lactams, co-trimoxazole, and Synercid. Cell division is the only process that upon interference, only generates increased sensitivity, specifically for CWSIs and co-trimoxazole. Interfering with peptidoglycan synthesis also mostly leads to increased sensitivity to CWSIs, as well as to 30S PSIs, while changes to genes that are involved in anchoring proteins to the cell wall (SP_1218 [*srtA*], SP_1833) can decrease sensitivity to CWSIs. Importantly, interfering with protein turnover, for instance through the protease complex ClpCP (SP_2194, SP_0746) and the regulator CtsR (SP_2195), which are generally assumed to be fundamental for responding to stress^44,45^, leads to decreased CWSI sensitivity and increased sensitivity to 30S and 50S PSIs (Fig. 3c and Supplementary Fig. 2). Moreover, FtsH (SP_0013), important for clean-up of misfolded proteins from the cell wall, increases sensitivity to 30S PSIs and Synercid, indicating how important protein turnover is especially for surviving 30S PSIs, which can trigger the production of faulty proteins. Most importantly, these data show that, as expected, hundreds of options exist where disruption of a pathway or process leads to increased sensitivity to specific antibiotics. Remarkably, there seem to be almost as many options that can lead to decreased antibiotic sensitivity.

**Figure 3.**
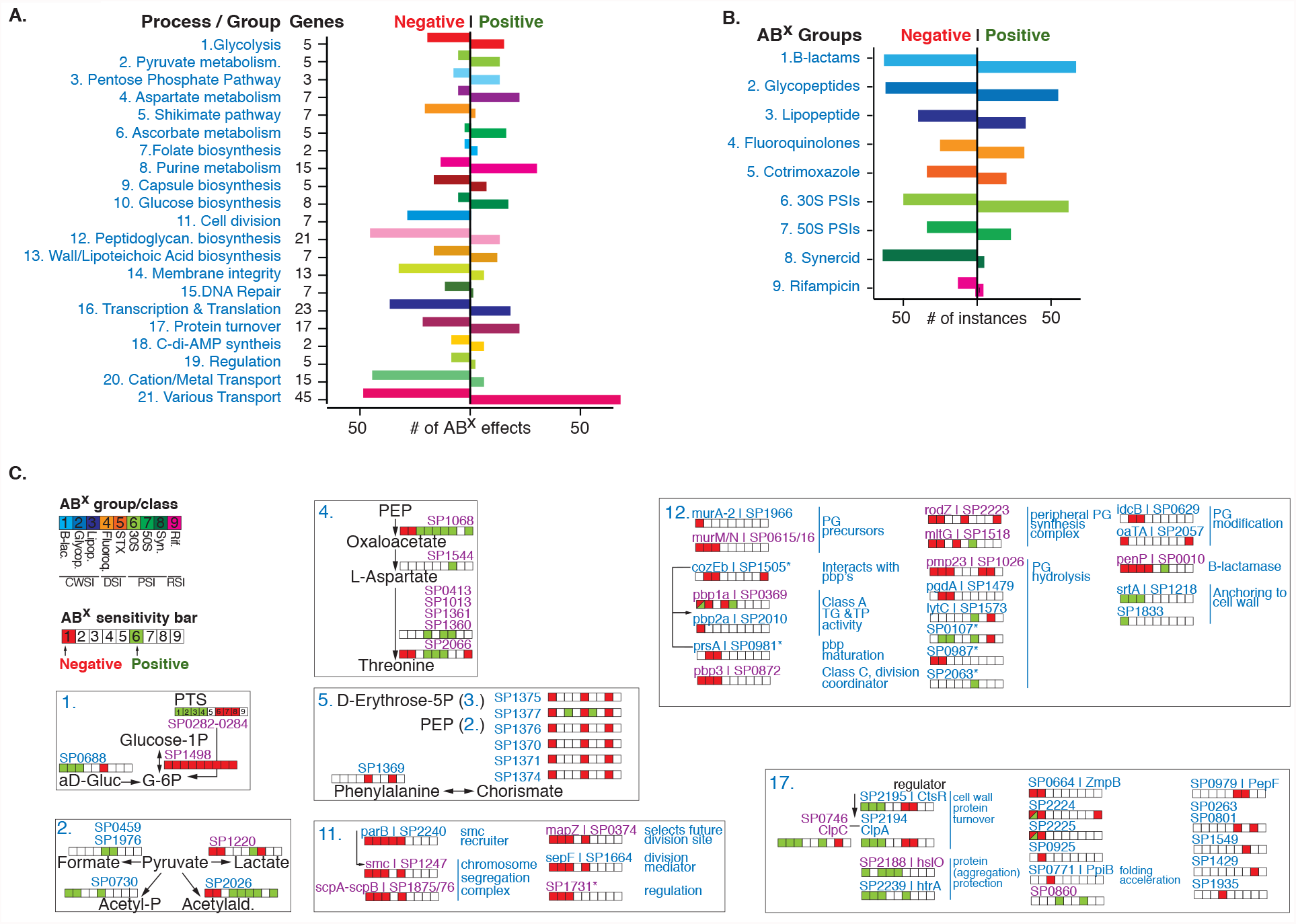
A multitude of options, pathways and processes can simultaneously lead to increased and decreased antibiotic susceptibility. **a**. Genes with at least one significant phenotype are split over 21 groups according to a pathway or process they belong to, which highlights how modulation of most pathways can lead to increased (negative) and decreased (positive) antibiotic sensitivity. **b**. While sensitivity to each antibiotic (group) can be increased by knocking out genes in the genome (negative), sensitivity can be decreased (positive) almost as often for most ABXs, except for Synercid, and to a lesser extent rifampicin, where most effects are negative. **c**. Detailed view of 7 out of 21 groups/processes highlighting how modulation of specific targets within each process leads to changes in antibiotic sensitivity. Each group is indicated with a number which is the same as in **a**. Where possible genes are ordered according to their place in a process/pathway, and gene numbers (SP_) are combined with gene names and annotation. Each indicated gene is combined with an ‘antibiotic sensitivity bar’ indicating whether disruption leads to increased (red/negative) or decreased (green/positive) sensitivity to a specific or group of antibiotics. When phenotypic responses are the same, multiple genes are indicated with a single bar (e.g. SP0282/SP0283/SP0284 in glycolysis, or SP0413/SP1013/SP1361/SP1360 in Aspartate metabolism). Gene numbers in blue have no effect on growth in the absence of antibiotics when knocked out, while gene numbers in purple have a significant growth defect in the absence of ABXs (see for detailed fitness in the absence and presence of antibiotics Supplementary Table 2). Essential genes are not indicated and genes with an asterisk have a partial or tentative annotation that has not been resolved. All 21 groups are listed in Supplementary Figures 2 and 3.

### *cozEb* encodes a cell division and peptidoglycan synthesis embedded membrane protein that can be critically targeted *in vivo* through an antibody-antibiotic strategy

By identifying targets that (re)sensitize bacteria against existing antibiotics, genome-wide antibiotic susceptibility data have the potential to guide the development of new antimicrobial strategies. One such strategy could be a combined therapeutic antibody-antibiotic approach; the antibody would target a gene-product that is important for sensitivity to one or more antibiotics and the product is easily accessible for the antibody at the bacterial cell surface. To find suitable candidate targets, Tn-Seq data were filtered for gene-products that, based on a known function or localization prediction, are likely to be present in the cell wall or membrane, and that when disrupted, increase sensitivity to one or more antibiotics. Moreover, it would likely be ideal if the gene is also important for survival *in vivo*. A strong candidate is SP_1505, which in the interaction network is most tightly linked to cell wall metabolism and cell division genes (Fig. 4a). After we previously hypothesized that it may play a role in cell wall integrity ^14^, it was recently named *cozEb*, with a likely role in organizing peptidoglycan synthesis during cell division ^46^, which fits its interaction profile (Fig. 4a). Importantly, the antibiotic Tn-Seq data suggest that disruption creates increased sensitivity to vancomycin and rifampicin, while the product is critical in the presence of daptomycin, which was confirmed through individual growth curves (Fig. 4b). The protein has eight predicted membrane-spanning domains (Fig. 4c), and *in vivo* Tn-Seq predicts it is important for survival in both the nasopharynx and lung (Fig. 4a, Supplementary Table 2). The gene was cloned into an expression plasmid generating an ∼30kD product (Fig. 4c), which was used to raise rabbit anti-CozEb antibodies, which were confirmed to be specific for the *cozEb* gene product (Fig. 4c). Potential antibody *in vitro* activity was determined through a bacterial survival assay in the absence and presence of antibodies and either vancomycin or daptomycin. Incubating bacteria with antibodies or daptomycin has no significant effect on bacterial survival, while vancomycin alone at the concentration used slightly reduces the number of surviving bacteria. Moreover, combining the antibody with either vancomycin or daptomycin further reduces the number of surviving bacteria *in vitro* compared to any agent individually (Fig. 4d). To assess whether the antibody-antibiotic approach works *in vivo*, mice were intranasally challenged with a bacterial inoculum either containing WT or *ΔcozEb*. Two additional sets of mice were challenged with WT and 8hrs post-infection they were either treated with daptomycin and control IgG-antibody or with daptomycin and CozEb-specific antibody. Mice were sacrificed 24 hrs post-infection, and bacteria in the lung were enumerated. As predicted by the *in vivo* Tn-Seq data the *cozEb* knockout has a significantly lower fitness in the lung highlighted by an up to 2.5-log lower bacterial load compared to WT. Importantly, while the WT survives equally well in the presence of the low daptomycin concentration and the control IgG antibody, in the presence of daptomycin and the CozEb-targeting antibody, its survival in the lung is significantly reduced and resembles that of the *cozEb* knockout (Fig. 4e). This shows that by combining antibiotic and *in vivo* Tn-Seq with gene annotation information, a gene-product can be selected that is central and critical to cell-wall synthesis and cell-division processes. Importantly, due to its presence in the membrane, it is directly targetable with an antibody, thereby sensitizing the bacterium to an antibiotic concentration it is normally not sensitive to.

**Figure 4.**
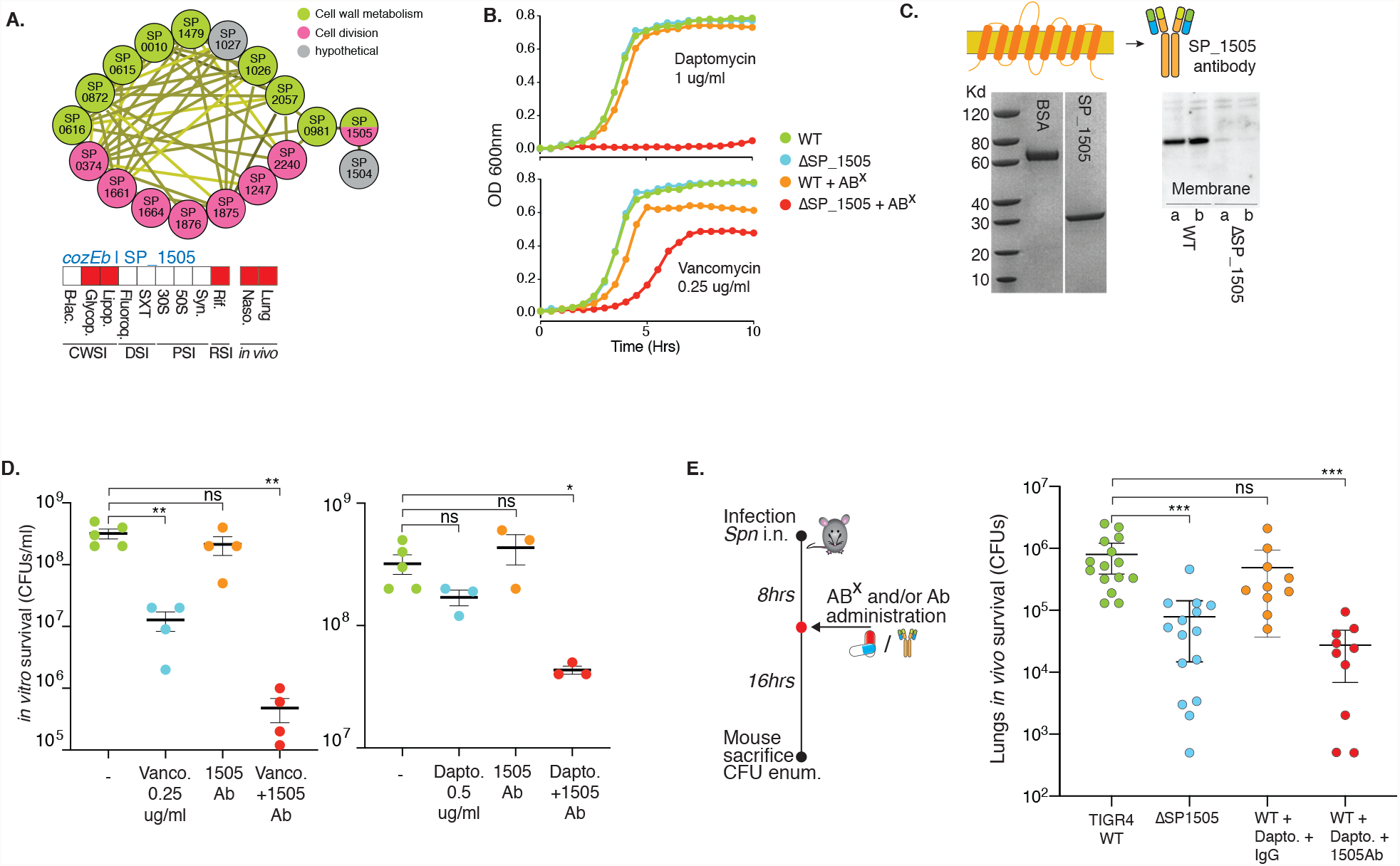
CozEb an integral membrane protein increases antibiotic sensitivity and can be targeted with an antibody. **a**. *cozEb*/SP_1505 is tightly clustered with cell division and cell wall metabolism genes, it is predicted to increase sensitivity to glycopeptides and the lipopeptide daptomycin, and has a decreased fitness in the mouse lung and nasopharynx. **b**. Growth curves of *ΔcozEb* validate its increased sensitivity to daptomycin and vancomycin. **c**. CozEb has 8 transmembrane domains, which generates a ∼30Kd product (BSA is shown as a control). The cloned protein was used to raise antibodies, which proofed to be specific for a product in the WT membrane, but does not bind anything in *ΔcozEb*, indicating the antibodies are specific for the membrane protein CozEb. **d**. Incubation of WT for 2hrs with vancomycin or daptomycin and in the presence of CozEb antibody, slightly but significantly decreases bacterial survival. **e**. An *in vivo* lung infection with WT or *ΔcozEb* confirms the mutant is less fit *in vivo*. While challenging the WT with daptomycin and IgG does not affect bacterial survival, a challenge with daptomycin and CozEb-specific antibodies, significantly reduces the recovered CFUs 24hrs post infection. Significance is measured through an ANOVA with Dunnett correction for multiple testing: *p<0.05, **p<0.01, ***p<0.001.

### The Ami-operon encodes an antibiotic importer, and inhibition triggers tolerance

While increased sensitivity profiles can guide the development of (re)sensitizing approaches, in contrast, the multitude of options that may lead to reduced antibiotic sensitivity (Fig. 3), could help in identifying (new) routes that may contribute to (the emergence of) antibiotic resistance. With lowered antibiotic sensitivity to 3 out of 4 antibiotic classes, the *ami*-operon is among genes with the greatest number of positive interactions. The operon forms a tight cluster in the interaction network (Fig. 3, 5a) and it is annotated as an oligopeptide transporter with no clear function. Two separate knockouts for SP_1888 *(amiE)* and SP_1890 *(amiC)* confirm decreased sensitivity to ciprofloxacin, vancomycin and gentamicin, and increased sensitivity to Synercid (Fig. 5b). There is limited evidence that the *ami*-transporter may have (some) affinity for at least two different peptides (P1 and P2) ^47-49^. These have been theorized to possibly function as signaling molecules and under certain circumstances may be generated by the bacterium itself ^47-49^. Both peptides were synthesized and while neither peptide affects growth of the WT or knockout mutants in the absence of antibiotics (Supplementary Fig. 4), the WT grows slightly better in the presence of gentamicin and peptide P2, but not P1 (Fig. 5b). This shows that some peptides may, at least partially, inhibit or occupy the *ami*-transporter, and thereby trigger decreased antibiotic sensitivity, in a similar manner as a knockout does. Besides peptides, the *ami-*transporter may be (non-selectively) transporting antibiotics into the cell, which could explain its effect on antibiotic sensitivity. To explore this, bacteria were exposed to ciprofloxacin or kanamycin and the internalized antibiotic concentration was determined through mass spectrometry for WT and both *ami* knockout mutants. In both mutants the amount of internalized ciprofloxacin was significantly lower (∼1.7x in *ΔamiE*, and ∼2.3x in *ΔamiC*), while the kanamycin concentration was found to be significantly lower in *ΔamiC* (∼2x; Fig. 5c). This shows that a functional *ami*-transporter increases the concentration of fluoroquinolones and 30S PSIs, suggestively by transporting them into the cell, and thereby, due to a higher internal concentration, enhancing the antibiotic’s inhibitory effects on growth. There are multiple examples that transporters can contribute to tolerance ^50,51^, which we recently showed is also the case for the *ade* transporter in *Acinetobacter baumannii*, which contributes to fluoroquinolone tolerance ^*7*^. However, those examples are mostly based on efflux pumps that actively decrease the antibiotic concentration in the cell through upregulation of such pumps. In contrast, with respect to the *ami*-operon it would be the reverse, i.e., inhibition instead of upregulation would lead to tolerance. To explore the effect on tolerance, the WT and *ΔamiE* were exposed to either 10xMIC of gentamicin or vancomycin over a period of 24hrs. Approximately 1% of the WT population survives 4hrs exposure to gentamicin, while none of the population survives exposure past 8 hrs. The *ΔamiE* population displays a slower decline in survival with 1% of the population surviving the first 8hrs (tolerant cells). At ∼10 hrs the decline ceases and the remaining population (∼0.01%) survives at least up to 24hrs, which is representative of a persister fraction ^21^. In contrast, the WT and *amiE* mutant populations decline at similar rates when exposed to vancomycin, showing that inhibition of the *ami*-transporter can lead to tolerance and persistence in an antibiotic specific manner.

**Figure 5.**
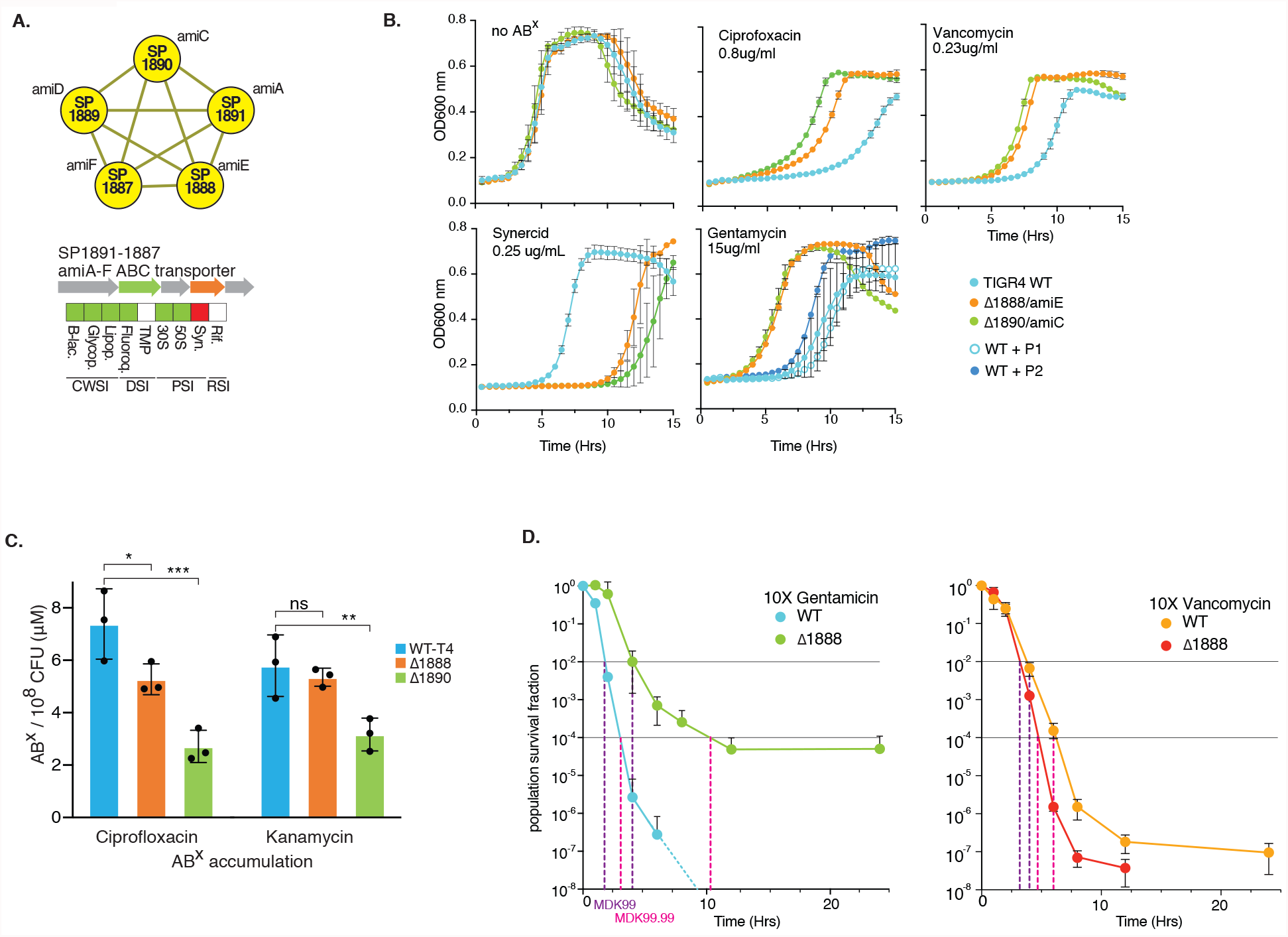
Modulation of the ami transporter decreases sensitivity to many antibiotics. **a**. The ami-operon forms a tight cluster, and upon knockout is predicted to decrease sensitivity to most antibiotics and increase sensitivity to Synercid. **b**. Growth curves of individual knockout mutants of *amiE* and *amiC* validate changes in antibiotic sensitivity and suggest the transporter phenotypically responds to peptide P2. **c**. Intracellular antibiotic accumulation analysis shows that the WT strain with an intact transporter reaches a higher intracellular antibiotic concentration, suggesting the transporter is involved in importing antibiotics, explaining why a knockout or occupation with a peptide such as P2, can lead to decreased antibiotic sensitivity. **d**. While modulation of the transporter leads to decreased sensitivity to gentamicin and vancomycin during growth, it leads to increased survival (i.e., tolerance) to gentamicin, but not vancomycin. Significance is measured through an ANOVA with Dunnett correction for multiple testing: *p<0.05, **p<0.01, ***p<0.001.

### Purine metabolism, (p)ppGpp and ATP production are tightly linked to altered ABX susceptibility and tolerance

Among the 21 functional groups, purine metabolism has some of the largest number of positive ABX interactions, mostly to β-lactams and glycopeptides (Fig. 3a, 6a). Moreover, two regulators (SP_1821/1979) associated with this pathway decrease sensitivity to β-lactams and/or glycopeptides and two ‘neighboring’ genes with unknown function have either the same (SP_0830), or the opposite effect (SP_1446) on antibiotic sensitivity as their defined neighbor, suggesting they may be involved in the same process as their neighbor (Fig. 6a). Furthermore, the global interaction network positively links an ABC-transporter (SP_0845-0848, Fig. 2c, 6a) with multiple genes in this pathway due to its similar profile. This operon is annotated as a putative deoxyribose-transporter, and to verify whether an interaction exists with purine metabolism, single and double knockouts were created between SP_0846 (the transporter’s ATP binding protein) and SP_0829/*deoB*. Their profiles suggest they do not affect growth in the absence of ABXs and have increased sensitivity to Synercid, which was confirmed in individual growth (Fig. 6b). However, when both knockouts are in the same background, their increased sensitivity to Synercid is masked. Thus, as indicated by the network, these results show that the ABC-transporter indeed has a genetic interaction with purine metabolism/salvage, but plays an unknown role. Importantly, this confirms that the global network includes valuable interactions that can be explored to uncover functional relationships.

**Figure 6.**
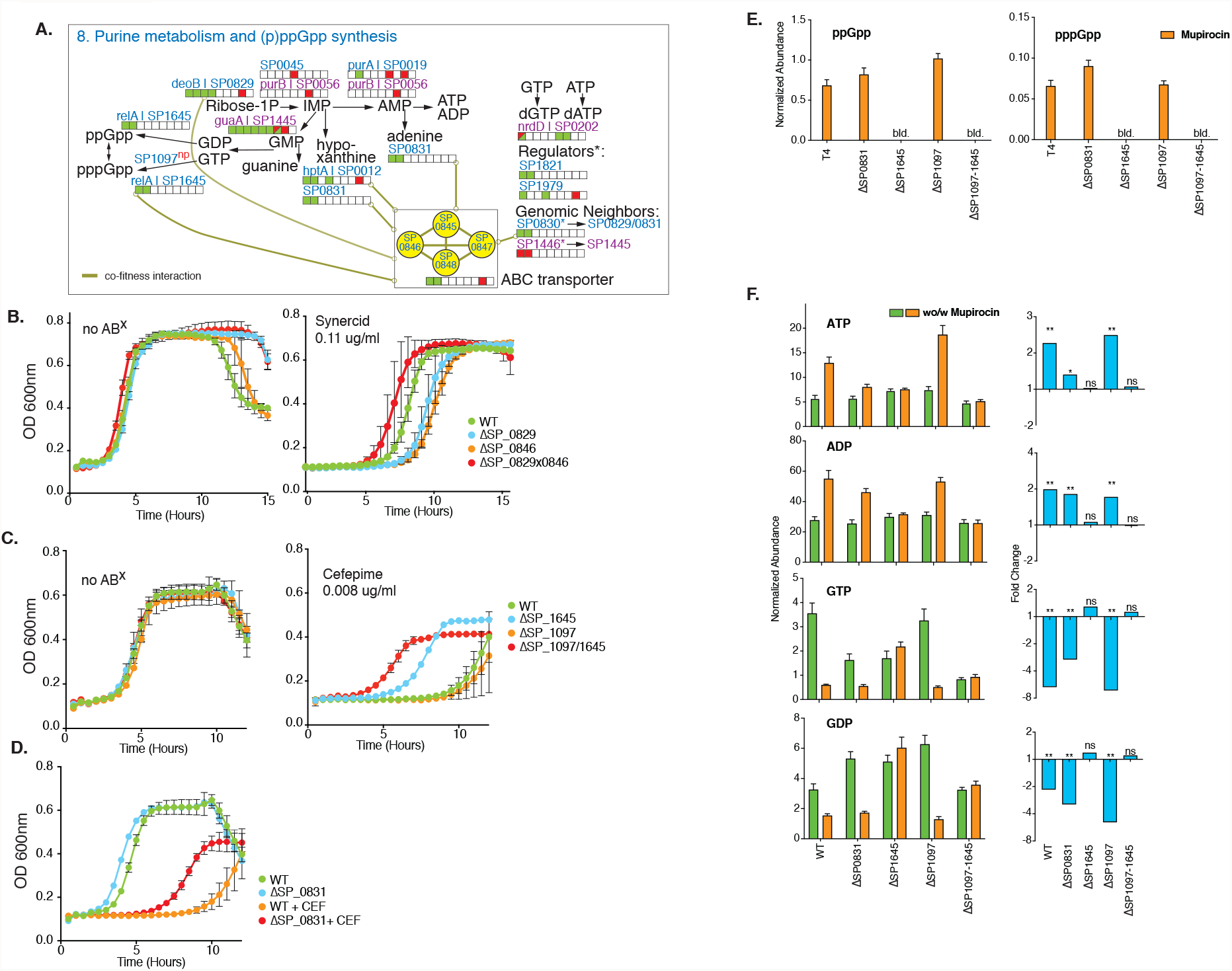
Modulation of purine metabolism affects alarmone and ATP synthesis and is linked to changes in ABX sensitivity. **a**. Several key steps of purine metabolism and their antibiotic sensitivity bars are indicated, with the same color coding as in Fig. 3. Note that for completeness SP_1097 is listed as well, for which we found no change in ABX sensitivity, which is denoted with ‘np’ for no phenotype. Also indicated is the putative deoxyribose transporter (SP_0845-0848) and its co-fitness interactions, which is shown as a high-connectivity cluster in Fig. 2. **b**. To determine whether the predicted interaction between SP_0845-0848 and purine metabolism leads to specific phenotypic changes, single knockouts were generated for deoB/SP_0829 and SP_0846, as well as a double knockout. While mutants and WT grow equally well in the absence of antibiotics, in the presence of Synercid, as predicted and indicated by their ABX sensitivity bar, the single knockouts display a higher sensitivity to the drug then the WT. The double mutant’s fitness in the presence of Synercid should change according to the multiplicative model if they act independently; i.e. their combined sensitivity should be the multiplicative of the individuals and thus further increase. Instead, the double knockout suppresses the increased sensitivity phenotype of the single mutants, indicating that the positive interaction that is found in the co-fitness network leads to a positive genetic interaction between these genes. **c**. Single and double knockouts of SP_1097 and SP_1645/*relA* grow just as well as WT in the absence of antibiotics. As predicted SP_1097 is equally sensitive to cefepime as the WT, while *ΔrelA* has decreased sensitivity as indicated by its ABX sensitivity bar in **a**. Additionally, the double knockout has decreased sensitivity to cefepime, indicating the dominant phenotype of *ΔrelA*. **d**. The phenotype of *Δ*SP_0831 was validated in growth as well, showing no change in growth in the absence of ABX, and decreased sensitivity in the presence of cefepime (FEP). **e**. The alarmone (p)ppGpp is below the limit of detection in the absence of stress (b.l.d.), upon induction with mupirocin it is synthesized in equal amounts in WT, *Δ*SP_0831 and *Δ*SP_1097, while it cannot be synthesized if *relA* is absent. **f**. Synthesis of di- and trinucleotides is significantly affected in the different mutants upon mupirocin exposure. Significance is measured through a paired t-test with an FDR adjusted p-value for multiple comparisons: *p<0.05, **p<0.01, ***p<0.001, ns not significant.

Furthermore, within purine metabolism the alarmone (p)ppGpp is synthesized from GTP and/or GDP. Like other bacterial species, *S. pneumoniae* likely responds to (some) ABXs via induction of the stringent response pathway^52^, in which *relA* (SP_1645) is the key player with both synthetase and hydrolase activity^53^. Additionally, SP_1097 is annotated as a GTP diphosphokinase and may be involved in the synthesis of pppGpp from GTP (Fig. 6a). Our data suggests, and we confirmed for the β-lactam cefepime (Fig. 6c), that when synthesis of the alarmone is inhibited by deletion of *relA*, similar to many other interactions in purine metabolism, this leads to reduced β-lactam and glycopeptide sensitivity (Fig. 6c). Moreover, while SP_1097, as predicted, does not change ABX sensitivity (Supplementary Table 2, Fig. 6), a double knockout of *relA*-SP_1097 seems to further decrease sensitivity to cefepime (Fig. 6c, Fig. 7a). Additionally, besides a change in growth, the single *relA* and double knockout (*ΔrelA*-SP_1097), also increases tolerance to cefepime by ∼1000-fold at 24hrs (Fig. 7b). To understand how *relA* and SP_1097 affect purine metabolism, we used LC/MS to measure (p)ppGpp, ADP, ATP, GDP and GTP. Additionally, we included SP_0831 a purine nucleoside phosphorylase involved in nucleotide salvage, which has the same ABX profile as *ΔrelA* (Fig. 6a, d), but should not directly affect (p)ppGpp synthesis. While (p)ppGpp is below the limit of detection during normal growth in any of the strains, as expected *ΔrelA* and the double mutant *ΔrelA*-SP_1097 are unable to synthesize the alarmone when exposed to mupirocin, a strong activator of the stringent response (Fig. 6e, Supplementary Table 9). In contrast, WT, *Δ*SP_0831 and *Δ*SP_1097 synthesize (p)ppGpp upon mupirocin exposure to a similar extent (Fig. 6e). Concerning the di- and trinucleotides in the pathway, upon mupirocin exposure GTP and GDP are significantly reduced in WT, *Δ*SP_0831 and *Δ*SP_1097, likely because they are used for (p)ppGpp synthesis (Fig. 6f, Supplementary Table 9). In contrast, while ATP and ADP again remain constant for the *ΔrelA* mutants, ATP and ADP synthesis are significantly increased upon mupirocin exposure, especially for WT and *Δ*SP_1097. This suggests that during activation of the stringent response, synthesis from IMP is directed towards AMP, and not necessarily GMP, at least not enough to replenish GTP and GDP. Additionally, upon mupirocin exposure, ATP only minimally increases for *Δ*SP_0831, while it increases over 2-fold for WT and *Δ*SP_1097 (Fig. 6f). It has been shown for bacteria including *Escherichia coli* and *Staphylococcus aureus* that a decreased ATP concentration can decrease sensitivity to ABXs such as ciprofloxacin^54^. Additionally, in *S. aureus* (p)ppGpp overexpression has been associated with decreased sensitivity to linezolid^55^. Our data suggests that (p)ppGpp and ATP synthesis may be intrinsically linked, i.e., at least in *S. pneumoniae* the inability to produce the alarmone also results in lowered ATP synthesis, which is associated with a lowered ABX sensitivity to β-lactams and glycopeptides. However, *Δ*SP_0831 shows that even if (p)ppGpp can be synthesized, modulation of purine metabolism, for instance through the salvage pathway, can result in decreased ATP synthesis, and can lead to lowered ABX sensitivity. Importantly, in many bacterial species, alarmone production is generally assumed to be triggered in response to different types of stress and has been shown to affect a large variety of processes including nucleotide synthesis, lipid metabolism and translation. (p)ppGpp is thereby a ubiquitous stress-signaling molecule that enables bacteria to generate a response that is geared towards overcoming the encountered stress. However, contradictory results between species indicates a possible non-uniformity across bacteria, leaving much to be learned about how the alarmone and the processes it can control fit into the entire organismal (response) network ^52^. Our data suggests that the inability (i.e., due to mutations) to generate the alarmone in *S. pneumoniae* in response to β-lactams and glycopeptides is linked to reduced ATP, which under specific circumstances may be an optimal response, as it results in decreased ABX sensitivity, and thereby a higher probability to survive the insult (Fig. 6c, 7a, b).

**Figure 7.**
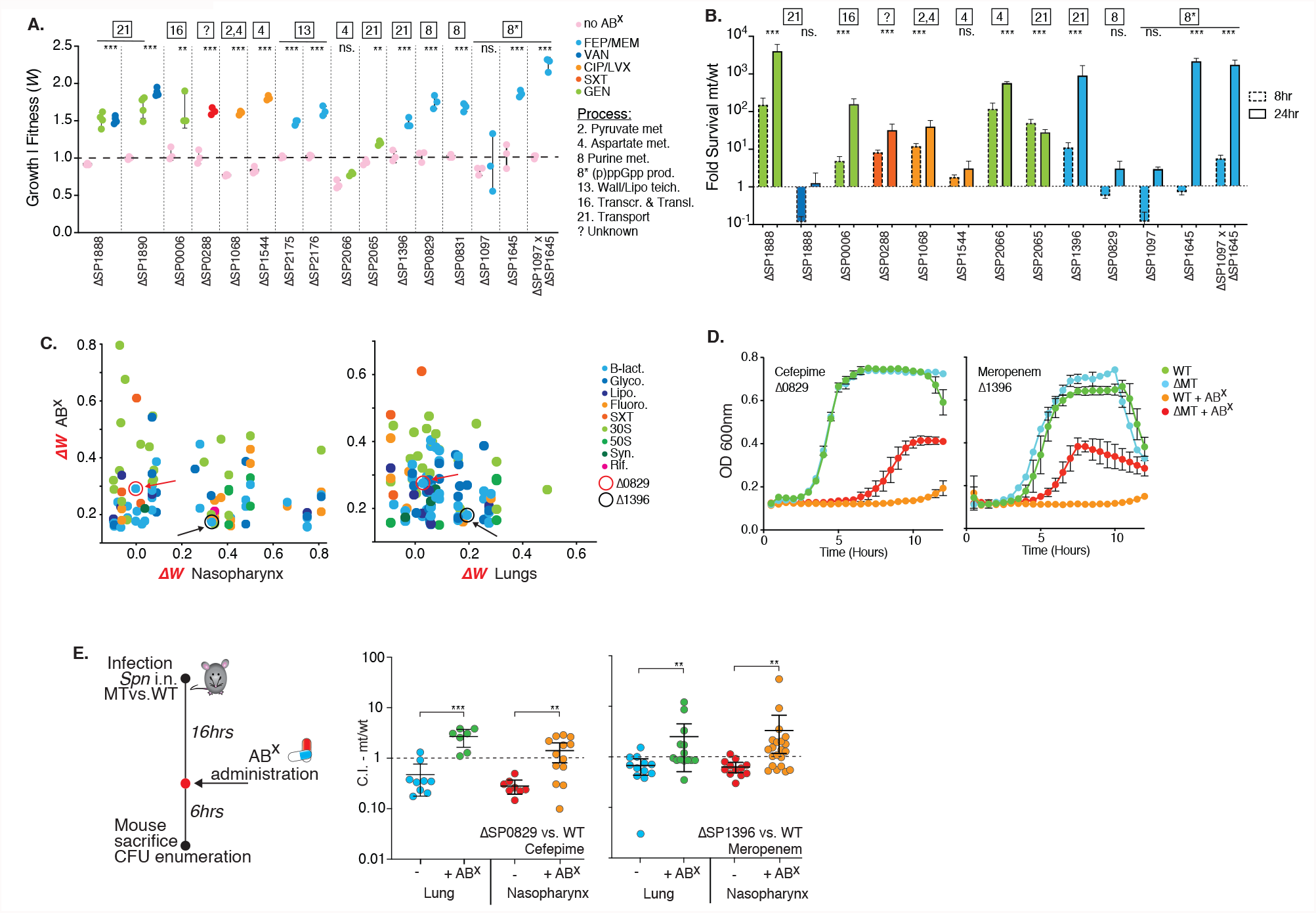
Decreased antibiotic sensitivity and tolerance can be achieved by modulation of a wide variety of processes. **a**. Relative growth rates of 16 knockout mutants involved in 7 processes measured in the presence of 7 antibiotics, validate that decreased ABX sensitivity can be achieved by modulating a wide variety of processes. **b**. Significantly increased survival during exposure to 5xMIC of an ABX over a 24hr period is observed for 9 out of 12 knockouts. Significance is measured with an ANOVA with Dunnett correction for multiple comparisons: **p<0.01, ***p<0.001. **c**. Tn-Seq data with a positive fitness in the presence of at least one antibiotic is plotted against *in vivo* Tn-Seq data showing those genes with only a small fitness defect, no defect or an increased predicted *in vivo* fitness, either during nasopharynx colonization or lung infection. Circled and indicated with arrows are SP_0829 in red and SP_1396 in black. **d**. *In vitro* growth curves validate decreased sensitivity to cefepime (SP_0829) and meropenem (SP_1396). **e**. Mice were challenged with WT and MT in a 1:1 ratio of which half received ABX 16hrs post infection (p.i.), and all were sacrificed 24hrs p.i. Displayed are the mutant’s competitive index (C.I.) in the nasopharynx and lung, and in the presence and absence of cefepime (SP_0829) or meropenem (SP_1396). In all instances, the addition of ABX significantly increases the C.I of the mutant. Significance is measured with a Mann-Whitney test **p<0.01, ***p<0.001.

### There are a multitude of predictable pathways that lead to tolerance *in vivo* in an antibiotic dependent manner

To further confirm that antibiotic sensitivity can be decreased by inhibiting a variety of processes, knockouts (KOs) were generated for fourteen mutants from 8 different processes. Thirteen mutants displayed an increased ability to grow in the presence of an ABX compared to the WT, and at least 8 mutants had an increased ability to survive high level exposure to an ABX (5-10xMIC) for at least 24 hours (Fig. 7a, b, Supplementary Table 8). Note that we validated 49 single KO genotype x phenotype associations in this study, with an equal distribution across the entire spectrum of ABX sensitivity (Fig. 1e, Supplementary Table 8). These data highlight that Tn-Seq data can be used to uncover a genome-wide ‘tolerome’, composed of a multitude of genes, pathways and processes that when modulated can decrease antibiotic sensitivity and/or trigger tolerance *in vitro* in an ABX dependent manner. Obviously, the selection regime *in vivo* is far more complex and stricter than in a test tube, which raises the question whether many of the *in vitro* tolerome options would be available *in vivo* as well. To explore this, all the Tn-Seq data with a positive fitness in the presence of at least one antibiotic was combined with *in vivo* Tn-Seq data and filtered for those genes with no or only a small fitness defect predicted *in vivo* during nasopharynx colonization or lung infection (Fig. 7c, Supplementary Table 2). Two genes were selected that we had confirmed for decreased ABX sensitivity *in vitro*: **1)** SP_0829/*deoB* synthesizes Ribose-1P and is involved in purine metabolism (Fig.6A). *ΔdeoB* has no effect on *in vitro* growth (Fig. 7a, d), it decreases sensitivity to cefepime during growth (Fig. 7a, d), but does not affect survival/tolerance (Fig. 7b); **2)** SP_1396/*pstA* is the ATP binding protein of a phosphate ABC transporter (Supplementary Fig. 3). *ΔpstA* has no effect on *in vitro* growth (Fig. 7a, d), it decreases sensitivity to meropenem during growth (Fig. 7a, d), and increases survival/tolerance (Fig. 7b). Both mutants were mixed with WT in a 1:1 ratio and used in an *in vivo* mouse infection competition model as we have done previously ^18^. Of the infected mice, half were administered antibiotics at 16hrs post infection, and were sacrificed 6hrs later to determine the strain’s competitive index (CI) (Fig. 7e). Importantly, while both mutants may have a slight disadvantage compared to the WT when colonizing the lung or nasopharynx, their CI increases significantly in the presence of ABXs, leading to increased survival compared to the WT (Fig. 7e, Supplementary Table 10). Combining antibiotic-with *in vivo* Tn-Seq thus confirms the existence of a wide-array of possible alterations of specific genes, pathways and processes that can have a beneficial effect *in vivo* in the presence of antibiotics. Such changes could thereby contribute to escape from antibiotic pressure and even create a path towards the emergence of antibiotic resistance.

With the possibility that some selective pressures in mice are similar in humans, this raises the possibility that stop codons in genes predicted by Tn-Seq to decrease antibiotic sensitivity while having no more than a minimal *in vivo* defect in the absence of ABXs, could be enriched for in antibiotic resistant clinical isolates. To test this hypothesis 4 gene-sets were compiled consisting of those that upon disruption: **1)** decrease antibiotic sensitivity in at least 1 antibiotic and have no strong defect *in vivo*; **2)** decrease antibiotic sensitivity in at least 1 antibiotic and have a defect *in vivo;* **3)** have little to no effect on antibiotic sensitivity and *in vivo;* **4)** have no effect or increase antibiotic sensitivity and have a defect *in vivo* (Fig. 8a, b; Supplementary Fig. 5). Thousands of strains were selected from the PATRIC^56,57^ database that could be split into a group of co-trimoxazole (SXT) resistant and a group of β-lactam resistant strains, and each group was matched with an equal number of sensitive strains from the database. In all strains in the SXT and β-lactam groups, irrespective of resistant or sensitive status, the number of stop codons in gene sets 1 and 3 are highest, which reflects the Tn-Seq predicted *in vivo* effects, i.e., while gene sets 1 and 3 contain mostly genes with potentially neutral effects, gene sets 2 and 4 contain many genes that are suggested to have a defect *in vivo* when disabled (e.g. with a stop codon) (Fig. 8c). Moreover, SXT resistant isolates in gene set 1 more often contain a stop codon compared to sensitive strains, and in β-lactam resistant isolates this is true for gene-sets 1-3 (Fig. 8d). While these are not ideal comparisons, for instance the entire ABX profile is not clear for many strains, different changes than premature stops could have ABX/*in vivo* modulating effects, strains could have experienced different ABX and/or *in vivo* selective pressures, and genetic changes can be strain-background dependent, it shows that genetic changes that can affect ABX and/or *in vivo* sensitivity, readily occur in clinical samples. This in turn underscores that ongoing infections may consist of variants that enable different paths to adjusting to, or overcoming a challenging host/ABX environment.

**Figure 8.**
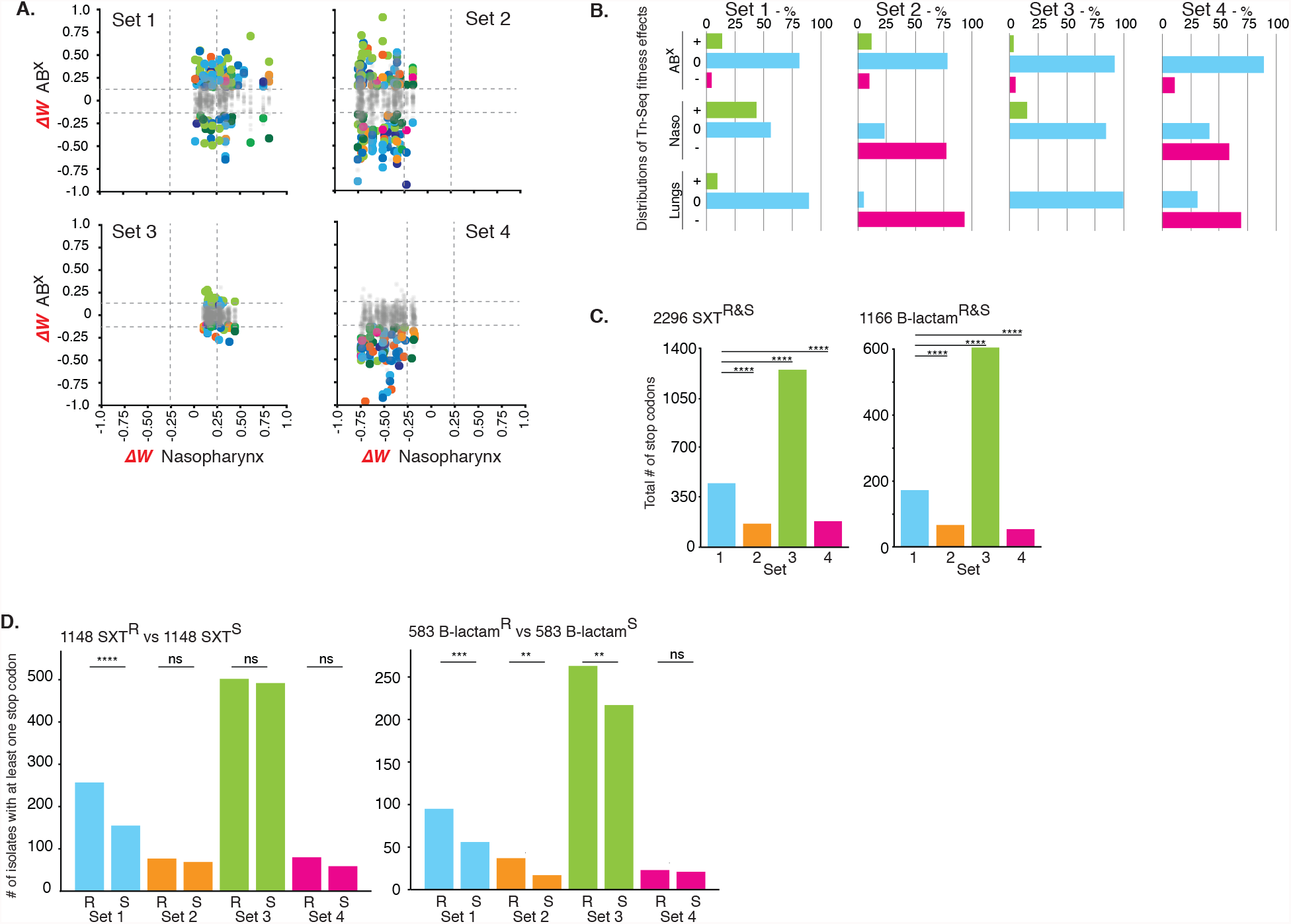
Stop codons are enriched in clinical samples in Tn-Seq predicted tolerome genes. **a**. Based on *in vivo* and ABX Tn-Seq data, four gene-sets consisting of 34 genes each were compiled with specific fitness profiles in the presence of antibiotics and *in vivo*. Shown are the *in vivo* effects for nasopharynx, while lung data are depicted in Supplementary Fig. 5. *ΔW* represents the fitness difference of a gene in a specific condition (e.g., an antibiotic, *in vivo)* minus its fitness *in vitro* in rich medium. Dashed lines indicate significance cut-offs, greyed-out dots indicate genes with no significant change in fitness in the presence of antibiotics, colors represent antibiotics and are the same as in Fig. 1. **b**. Detailed distributions for each gene set highlights whether effects in the presence of antibiotics, in the nasopharynx and lungs increase (+), do not affect (0) or decrease (-) relative fitness. Gene set rationales are described in the text. **c**. The total number of stop codons in each gene set for 2296 co-trimoxazole and 1166 β-lactam resistant and sensitive strains. **d**. The number of sensitive and resistant strains with at least one stop codon in a gene in each gene-set. Significance is measured through a Fisher’s exact test: **p<0.01, ***p<0.001, ****p<0.0001.

## CONCLUSION

The emergence and increase in antibiotic resistance among most bacterial pathogens is a continuously developing problem with several important drivers, which include: **1)** a lagging development of new drugs and treatment strategies; **2)** a lack of (rapid) diagnostics and prognostics; and **3)** an incomplete understanding of how antibiotic resistance develops. Moreover, these drivers are inherently connected making it a complex problem to solve. First, the ability of bacteria to evolve resistance elicits an arms-race that requires the development of new drugs and treatment strategies to keep the balance of infection-control tipped in our favor. Thus, while developing new drugs would keep the arms-race in place, the ability to slow or prevent the emergence of resistance could resolve the status quo. Furthermore, even though it is critical to understand how and under which circumstances resistance evolves, the applicability of this knowledge depends on the availability of diagnostics that could inform on the emergence of resistance (precursors) and thereby guide and enable timely, tailored and targeted treatments. To progress towards a comprehensive understanding of how an infection is developing in the absence or presence of treatment, and how to decide what to do next, we believe that a detailed genetic understanding of how a bacterium deals with and overcomes stress, as well as its genetic potential to achieve this, are key aspects. In this study we contribute to reaching such an understanding by building and exploring a detailed atlas of ABX sensitivities, which highlights how modulation of specific genes, pathways and processes does not only result (as expected) in increased ABX sensitivity, but almost just as often in decreased ABX sensitivity. We show that such an atlas can be used to identify leads for gene function, to uncover the genome’s underlying architecture and genetic relationships among genes, for the identification of new drug targets, and the development of new proof-of-principle antimicrobial (ABX sensitizing) strategies. Most importantly, these data identify genome-wide genetic changes that show how modulation of genes, pathways and processes can lead to lowered antibiotic sensitivity and tolerance, not only *in vitro*, but also *in vivo*. Moreover, we show that mutations that have the potential to trigger the same phenotypes readily occur in patients. These detailed data on reduced antibiotic sensitivities thereby suggest that far more potential routes to ABX-escape, and potentially resistance, may exist than assumed. However, it does not exclude that (multiple) general mechanisms exist that can trigger such decreased sensitivities. For instance, the overlap in ABX sensitivity profiles (e.g. decreased sensitivity to CWSIs) that emerge from modulating specific parts of glycolysis, pyruvate, ascorbate, glucose and purine metabolism, protein turnover and (p)ppGpp and c-di-AMP synthesis (Supplementary Fig. 2 and 3), could possibly all be linked by a common effect, that may at least partially come from a decreased ATP availability. Moreover, while a slower growth rate is also often linked to decreased ABX sensitivity and tolerance, we show that it is not the driving force behind our results as most of the created KOs have no effect on growth in the absence of ABXs. Importantly, we believe these data are both an argument and potential starting point for a platform to predict clinically relevant mutations and determinants of antibiotic resistance/tolerance. Consequently, these results underscore the importance of understanding the genetics of variants with altered drug susceptibility, as their genetics makes them diagnostically identifiable and trackable, while their often-associated collateral sensitivities to other ABXs or drugs could make them targetable.

## METHODS

### Bacterial culturing, growth curves and tolerance experiments

Experiments were performed with *S. pneumoniae* strain TIGR4 (NCBI Reference Sequence: NC_003028.3). TIGR4 is a serotype 4 strain that was originally isolated from a patient from Norway with Invasive Pneumococcal Disease (IPD) ^58,59^. All ‘SP_’ gene numbers in the tables and figures are according to the TIGR4 genome. Single gene knockouts were constructed by replacing the coding sequence with a chloramphenicol and/or spectinomycin resistance cassette as described previously ^18,35,36^. *S. pneumoniae* was grown on sheep’s blood agar plates or statically in THY, C+Y or semi-defined minimal media at pH 7.3, with 5 μl/ml Oxyrase (Oxyrase, Inc), at 37°C in a 5% CO_2_ atmosphere ^15^. Where appropriate, cultures and blood plates contained 4 μg/ml chloramphenicol (Cm) and/or 200 μg/ml spectinomycin (Spec). Single strain growth assays were performed three times using 96-well plates by taking OD_600_ measurements on a Tecan Infinite 200 PRO plate reader or BioSpa 8 (BioTek). Tolerance experiments were performed by exposing exponentially growing bacteria to ∼10xMIC of an antibiotic. Samples were taken at different time-points over a 24hr period, washed with PBS and plated on blood-agar for enumeration.

### Tn-Seq experiments, fitness (*W)* and enrichment analyses

Six independent transposon libraries, each containing ∼10,000 insertion mutants, were constructed with transposon Magellan6 in WT-T4 as described previously ^14,18,35,60^. Selection experiments were conducted in rich medium with glucose as a carbon source in the presence or absence of 20 different antibiotics at a concentration that slows growth by ∼30-50% (Supplementary Table 1). Sample preparation, Illumina sequencing and fitness calculations were done as described ^14,18,35,40,60,61^. In short, for each insertion, fitness *W*_*i*_, is calculated by comparing the fold expansion of the mutant relative to the rest of the population by using an equation that we specifically developed to have fitness represent the growth rate of a mutant^18,35,61^. All of the insertions in a specified region or gene are then used to calculate the average fitness and standard deviation of the gene knockout in question. This means that *W*_*i*_ represents the growth rate per generation, which makes fitness independent of time and enables comparisons between conditions. To determine whether fitness effects are significantly different between conditions three requirements have to be fulfilled: 1) *W*_*i*_ is calculated from at least three data points, 2) the difference in fitness between the presence and absence of antibiotic has to be larger than 15% (thus *W*_*i*_ - *W*_*j*_ = < −0.15 or > 0.15), and 3) the difference in fitness has to be significantly different in a one sample *t*-test with Bonferroni correction for multiple testing^18,35,61^. To determine whether a particular process or pathway is specifically involved in responding to an antibiotic-group, a hypergeometric test was performed to test for enrichment. The distribution of significant genes within each process was compared to the distribution of the pathways in the overall genome. A p-value and Benjamini-Hochberg adjusted p-value were calculated for each process and antibiotic group, where an adjusted p-value below 5% is considered to identify statistical enrichment.

### Co-fitness network construction and SAFE analysis

A gene x condition matrix was constructed to identify correlating fitness profiles and built a co-fitness network. The matrix is based on 20 antibiotic conditions from experiments performed here, supplemented with 17 conditions from van Opijnen and Camilli 2012^18^ (Supplementary Table 3). The additional conditions consist of Sucrose, Fructose, Cellobiose, Raffinose, Sialic Acid, Galactose, Mannose, Maltose, GlcNac, Bipyridyl, transformation, hydrogen-peroxide, methyl-methane sulfonate, pH6, temperature, Norfloxacin. Genes with missing data were removed resulting in a 1519 gene x 37 condition matrix (Supplementary Table 3). Genes and conditions were correlated using a Pearson’s correlation coefficient and a Spearman’s correlation coefficient. Resulting in two 1519×1519, gene vs gene matrices. A significance cutoff was applied and correlations ≥0.75 were retained and used as edges to build a co-fitness network consisting of 1519 genes and 2399 edges. An edge-weighted spring embedded layout was applied with Cytoscape ^62^, with the absolute correlation value as the edge weight. This results in a network with several major clusters and multiple genes unconnected to the main network. A stability test was performed to determine the robustness and quality of each edge in the network by building a correlation matrix from partial data. 30 conditions were selected 100 times to build a correlation matrix and using the same cutoff criteria a co-fitness matrix was compiled. Every edge with a correlation value above the threshold was assigned a 1 and every edge below the cut-off 0. This resulted in 100 binary matrices which were then summated, resulting in every gene vs gene interaction being assigned a stability score with a value N out of 100. A SAFE (Spatial Analysis of Functional Enrichment) analysis^41,42^ on the co-fitness network was performed with Cytoscape. A SAFE analysis is geared towards defining local neighborhoods for each node within a network and calculates an enrichment score for every functional attribute. It then highlights the areas that are the most enriched for that attribute. Attributes were assigned by merging KEGG^63^ pathway annotation and available functional category annotations, which covers 94% of the genes within the network. The distance threshold was set to the 1^st^ percentile of the map-weighted distance, the Jaccard similarity index was set to 0.5, and nodes in different landscapes were retained.

### CozEb (SP_1505) cloning and protein expression

Cloning and expression of SP_1505 was undertaken commercially (Genscript). Codon-optimized SP_1505 was cloned into pET28a with a C-terminal His-tag. *E. coli* BL21 (DE3) was transformed with recombinant plasmid. A single colony was inoculated into LB medium containing kanamycin; cultures were incubated at 37°C at 200 rpm and IPTG was introduced for induction. SDS-PAGE and Western blot were used to monitor the expression. Protein was purified from 1L batch culture in Terrific Broth. Cells were harvested by centrifugation, cell pellets were lysed by sonication, and supernatant after centrifugation was kept for future purifications. SP_1505 protein was obtained by three-step purification using Ni column, Superdex 200 column and Q Sepharose. Fractions were pooled and dialyzed followed by 0.22 μm filter sterilization. Protein was initially analyzed by SDS-PAGE and Western blot by using standard protocols for molecular weight and purity measurements. The primary antibody for Western blot is Mouse-anti-His mAb (GenScript, Cat.No.A00186). The concentration was determined by BCA protein assay with BSA as a standard. Final protein product was stored in 50 mM Tris-HCl, 150 mM NaCl, 10% Glycerol, 0.2% DDM, pH 8.0 and stored at −80°C.

### CozEb (SP_1505) antibody generation, purification and quantification

A single rabbit was vaccinated by a commercial vendor (Rockland) with recombinant SP_1505 via the following schedule. Rabbit was immunized via intradermal route with 0.1 mgs SP_1505 with Complete Freund’s Adjuvant (CFA) followed by an intradermal 0.1 mg booster injection with Incomplete Freund’s Adjuvant IFA as an adjuvant at day 7, followed by two subcutaneous 0.1 mg booster injections at days 14 and 28 with IFA. Terminal bleed was collected on day 52 following challenge. SP_1505 IgG was purified from immunized rabbit serum using protein G resin and columns (Pierce) according to manufacturer specifications. Following purification, antibody was concentrated using 10,000 MWCO centrifugal filters (Millipore) and was dialyzed three times against PBS in a 3.5kDa Slide-A-Lyzer dialysis cassette (Thermo Scientific). Antibody specificity was determined by Western Blot using the parental wild-type and corresponding deletion mutants.

### Cell fractionation, TCA precipitation and Western Blotting

Strains were grown in Todd-Hewitt broth to OD 0.4. Following this, cells were fractionated as previously described^64^. Briefly, 2mL of culture was centrifuged at maximum speed. The pellet was resuspended in cell wall digestion buffer [1x Protease inhibitor cocktail (Roche), 300U/uL mutanolysin, 1mg/mL lysozyme in a 30% sucrose-10mM Tris (pH 7.5) buffer with 20mM MgCl2 and 20mM MES (pH 6.5)] and incubated at 37°C for 60 minutes. After centrifugation, the supernatant containing the cell wall was saved. Pelleted protoplasts were snap frozen in a dry ice ethanol bath, then treated with MgCl2, CaCl2, DNase I (Qiagen), and RNAse A (Roche) in 50mM Tris buffer (pH 7.5) with 20mM HEPES (pH 8.0), 20mM NaCl, and 1mM DTT with protease inhibitors. The pellet was incubated on ice for one hour, then spun at max speed for 30 minutes at 4°C. The supernatant, which contained the cytoplasmic fraction, and the pellet, which contained the membrane fraction, were saved. 100% TCA was added to the samples so that the final concentration of TCA was 20%. Samples were incubated on ice for 30 min, then centrifuged at full speed at 4°C to pellet precipitated protein. The TCA supernatant was aspirated, and the pellet was washed twice with 100% acetone, then air-dried at 95°C for 1 minute. Pellets were resuspended in NuPage LDS sample buffer (Thermo Scientific) and boiled at 100°C for 10 minutes. Samples were loaded into NuPage SDS-PAGE gels (Thermo Scientific) and transferred to nitrocellulose membranes using the XCell Sure-Lock mini-cell electrophoresis system (Thermo Scientific). Nitrocellulose membranes were blocked overnight in 5% NFDM and treated with primary antibody against SP_1505 at a concentration of 1:500. After washing, membranes were treated with secondary antibody goat anti-rabbit IgG-HRP (Bio-Rad) at a concentration of 1:3000. Membranes were developed using the SuperSignal West Dura Extended Duration Substrate (Thermo Scientific) and were visualized using a BioRad ChemiDoc MP imaging system.

### Antibiotic-antibody targeted *in vitro* bacterial survival

Bacteria were inoculated from TSA plates into C+Y media, at OD 0.4, culture was split into 1 mL aliquots and treated with vancomycin (0.25 μg/ml) or daptomycin (0.5 μg/ml). For antibody treatment, strains were grown in C+Y media until OD 0.3. At this time, samples were treated with SP_1505 antibody or control rabbit IgG antibody (Sigma) at concentrations indicated in figure legends, incubated for 30 minutes, followed by antibiotic treatment. At 4hrs post antibiotic addition samples were plated for bacterial enumeration.

### Antibiotic-antibody mouse challenge

Isoflurane-anesthetized 7-week-old female BALB/c mice were inoculated intranasally with 10^6^ CFU of wild type pneumococcal cells in a volume of 100 μL. Eight hours following the challenge mice were treated with vehicle (Plasmalyte), vancomycin (0.25 mg/kg), daptomycin (2.5mg/kg), a-SP_1505 antibody (100 uL), and control rabbit IgG. At 16 hours following antibody/antibiotic treatment (24hr post-challenge) mice were euthanized, and lungs and chest cavity blood were removed for quantification of bacteria. Whole lungs were washed twice in PBS, and lung tissue was subsequently homogenized in 1 mL PBS. Homogenized lung samples were centrifuged at 300xg, and bacteria-containing supernatant was plated onto Neomycin-containing blood agar plates for CFU titers.

### Peptide production

Peptide P1 (Ser-Asn-Gly-Leu-Asp-Val-Gly-Lys-Ala-Asp) and peptide P2 (Ala-Lys-Thr-Ile-Lys-Ile-Thr-Gln-Thr-Arg) were synthesized on a preloaded Wang resin using the standard Fmoc/tBu chemistry for peptide synthesis. All coupling reactions were carried out in DMF using HBTU as the coupling reagent, 0.4 N-Methyl Morpholine in DMF as base. After each coupling, deprotection of the Fmoc group was done by using 20% piperidine in DMF. After completion of synthesis, peptides were cleaved from resin using TFA and purified using RP-HPLC. The integrity and purity of the peptides were confirmed using LC-MS.

### Antibiotic accumulation

Antibiotic accumulation was determined as previously described^65^. *S. pneumoniae* were grown in THY to OD 0.6. Cells were pelleted, washed twice in PBS and resuspended in 3.5mL PBS. 1mL of cells were incubated with 50μM antibiotic for 10 minutes at 37^°^C. Following incubation, 800μL of drugged cells were spun (3min, 13,000xg) through 700μL of a 9:1 mix of AR20 and high temperature silicon oils (cooled to - 80 ^°^C), after which the supernatant of silicone oil and free compound were carefully removed. For lysis, pelleted cells were resuspended in 200μL dH_2_O and lysed via bead beating (3x 15s at 5m/s). Debris was pelleted (10’ at 20,000xg) and 100μL of supernatant was removed and saved. Cell debris was resuspended in the remaining 50μL dH_2_O and mixed with 200μL methanol. Potential cell debris was pellet again and 150μL of the methanol extract was mixed with the 200μL dH_2_O supernatant from the previous step. The extract was pelleted one final time (10’ at 20,000g) before being filtered (0.22μm).

Samples were analyzed with a Waters Acquity M Class series UPLC system and Xevo G2 QTOF tandem MS/MS with Zspray. 100nl of extract was separated using a Phenomenex Kinetex 2.6 μm XB-C18, 100 Å (300 μm × 150 mm) column with solvent A, 0.1% formic acid in water, and solvent B, 0.1% formic acid in acetonitrile. The inlet method utilized a flow rate of 8 μl min^−1^ with the following gradient: 0−4 min, 99.9% solvent A and 0.1% solvent B; 4–5 min, 10% solvent A and 90% solvent B; 5–6 min, 99.9% solvent A and 0.1% solvent B. Tandem mass spectra were acquired with the following conditions: Ciprofloxacin: CV:20, CE:25, m/z ion: 333.14.→245.11; Kanamycin: CV:40, CE:20, m/z ion: 485.25→163.11. High-resolution spectra were calibrated by co-infusion of 2 ng ml^−1^ leucine enkephalin lockspray (Waters). Data were quantified using Waters MassLynx software where the AUC was determined by integrating the corresponding daughter peak of the parent compound. Concentrations of the unknown compounds were determined by the linear fit of the corresponding standard curve. Concentrations are reported as the average of three biological replicates.

### (p)ppGpp induction and LC/MS analysis

*S. pneumoniae* strains were grown at 37 °C in 10 mL ThyB to an OD of ∼0.5. Cultures were split into 5 mL aliquots for mupirocin-treated versus untreated controls. To induce the stringent response and ppGpp production, mupirocin was added in a final concentration of 25 μg/mL and incubated at 37 °C for 30 minutes. Cells were centrifuged at 6000× g for 5’, supernatant was discarded and cell pellets were frozen at −80 °C. For LC/MS analysis cell pellets were resuspended in 2 ml cold methanol, and 150 pmol of [^13^C_10_]-GTP (Sigma) was added and incubated at −80 °C for 30 minutes. Samples were centrifuged at 4000 x g for 10’, and the supernatant was removed and dried overnight in a Savant Speedvac Concentrator SPD 1010 (Thermo Fisher). Samples were analyzed using a Shimadzu Prominence UFLC attached to a QTrap 4500 equipped with a Turbo V ion source (Sciex). Samples (5 μL) were injected onto a SeQuant ZIC-cHILIC, 3 μm, 2.1 × 150 mm column at 30 °C (Millipore) using a flow rate of 0.3 ml/min. Solvent A was 25 mM ammonium acetate, and Solvent B was 75% acetonitrile + 25 mM ammonium acetate. The HPLC program was the following: starting solvent mixture of 0% A/100% B, 0 to 2 min isocratic with 100% B; 2 to 4 min linear gradient to 85% B; 4 to 17 min linear gradient to 65% B; 17 to 22 min isocratic with 65% B; 22 to 25 min linear gradient to 100% B; 25 to 30 min isocratic with 100% B. The QTrap 4500 was operated in the negative mode, and the ion source parameters were: ion spray voltage, −4500 V; curtain gas, 30 psi; temperature, 400 °C; collision gas, medium; ion source gas 1, 20 psi; ion source gas 2, 35 psi; declustering potential, −40 V; and collision energy, −40 V. The MRM transitions are: ppGpp, 602.0/159.0; pppGpp, 682.0/159.0, and [^13^C_10_]-GTP, 522.0/159.0. [^13^C_10_]-GTP was used as the internal standard. The system was controlled by the Analyst software (Sciex) and analyzed with MultiQuant™ 3.0.2 software (Sciex). Peaks corresponding to ppGpp and pppGpp were quantified relative to the internal standard. The limit of detection for ppGpp and pppGpp is 5 pmol, and for GTP, GDP, ATP and ADP 0.05pmol.

### *In vivo* mouse competition experiment wo/w antibiotics

*In vivo* competition experiments were essentially performed as previously described (van Opijnen and Camilli 2012). Specifically, groups of at least 12 outbred 4-6-week-old Swiss Webster mice (Taconic Inc.,) were anesthetized by isoflurane inhalation and challenged intranasally (i.n.) with 50 μl, ∼1.5 × 10^7^ CFU, bacterial suspension in 1X PBS. Each bacterial suspension contained a 1:1 mixture of *S. pneumoniae* TIGR4 wildtype and ΔSP_0829 or ΔSP_1396. The challenge dose was always confirmed by serial dilution and plating on blood agar plates. Infected mice receiving antibiotic treatment were administered either 1 mg/kg cefepime (WTvsΔSP_0829) or 10 mg/kg meropenem (WTvsΔSP_1396) 16 hours post-bacterial challenge by intraperitoneal (i.p.) injection. Antibiotic dosing was previously determined to reduce bacterial loads 10-100-fold *in vivo*. Mice were euthanized by CO_2_ asphyxiation at 6 hours post-antibiotic administration (or 22 hours post-bacterial challenge). Blood by cardiac puncture, nasopharynx lavage, and total homogenized lungs were collected from each animal to determine bacterial burden by serial dilution and plating blood agar plates as previously described^18^.

### Clinical-strain stop-codon analysis

Four gene-sets were compiled to test for the differential occurrence of stop-codons in patient samples. Each gene-set consists of 34 genes and are defined as: **Set 1** consists of genes that when disrupted lead to a significant decrease in antibiotic sensitivity in the presence of at least one antibiotic (*in vitro* ABx fitness positive), and have no fitness defect in lung and nasopharynx (*in vivo* neutral or positive); **Set 2** consists of genes that when disrupted lead to a significant decrease in antibiotic sensitivity in the presence of at least one antibiotic (*in vitro* ABx fitness positive), and have a significant fitness defect in lung and nasopharynx (*in vivo* fitness negative); **Set 3** consists of genes that when disrupted have no fitness benefit in any of the antibiotics (*in vitro* ABx fitness neutral), but with a significant fitness benefit in lung and nasopharynx (*in vivo* fitness positive); **Set 4** consists of genes that have decreased fitness in the presence of antibiotics (*in vitro* ABx fitness negative), and that have a significant fitness defect of >15% in lung and nasopharynx (*in vivo* fitness negative). The PATRIC database was screened for antibiotic resistant *S. pneumoniae* isolates. There is a potential risk that isolates in the database are clonally related, which could mean that multiple isolates would contain exactly the same sequence and for instance the same stop codon, which could bias the analysis. To reduce this potential bias candidate isolates were limited to those belonging to a different MLST type. While this considerably reduced the number of potential isolates, we were able to collect 533 β-lactam resistant and 1147 co-trimoxazole resistant strains. Moreover, an equal number of non-resistant strains were compiled. From each genome, gene sequences were extracted that match those from each of the 4 gene-sets. Each gene was scanned for premature stop codons occurring in the first 90% of a gene. For each gene-set the number of strains with at least one stop codon in the gene-set were recorded, as well as the total number of stop-codons in all genes in a set. To test for differences in the number of isolates containing a stop codon within (susceptible vs. resistant) and between sets a Fisher’s exact test was performed.

## ACKNOWLEDGEMENTS

DNA sequencing was performed at the Boston College Sequencing Core. The authors wish to thank Jon Anthony for running the *Aerobio* sequencing analyses pipeline, and Ralph Isberg and Vaughn Cooper for valuable discussions. This work was supported by a Charles King Trust Fellowship to F.R., NIH R01 AI110724 to T.v.O., and U01 AI124302 to T.v.O. and J.W.R.

## AUTHOR CONTRIBUTIONS

T.v.O. devised the study and wrote the manuscript. E.R, B.S, L.M.N.R, F.R, A.N, S.J.W, B.J, N.B, K.L, J.G, M.F, S.M.R, R.E.L, C.R, J.W.R, and T.v.O. performed wet-lab experiments, data collection and interpretation. D.L. and T.v.O. performed Tn-Seq data analysis, pathway and network construction, analysis and interpretation. J.W.R. contributed to key conceptual ideas. All authors contributed to manuscript editing and approved the final paper.

## DATA AVAILABILITY

Sequencing data is available at the Short Read Archive (BioProject accession number PRJNA750080).

